# High-resolution NMR spectroscopy of proteins in intact mitochondria

**DOI:** 10.1101/2025.03.19.644076

**Authors:** Zeting zhang, Cai zhang, Guohua Xu, Ruichen Du, Jinbo Yu, Xiaoli Liu, Zhaofei Chai, Qiong Wu, Ling Jiang, Maili Liu, Conggang Li

## Abstract

Understanding protein structure and function within mitochondria is essential for unraveling the molecular mechanisms underlying cellular energy production, stress response, and disease. Here, we present a novel approach for NMR observation of proteins within intact mitochondria by delivering proteins directly into isolated mitochondria via electroporation. Using this method, we investigate the interaction of α-synuclein (α-syn) with the mitochondria membrane and examine how post-translational modifications regulate this interaction. Additionally, we assessed the stability of GB1 and the dimerization of its variant within mitochondria, achieving quantitative insights into mitochondrial environmental impact on protein function. This approach offers a valuable framework for exploring mitochondrial related biomolecular events at atomic-resolution within intact mitochondria, paving the way for a more comprehensive understanding of the molecular events governing mitochondrial health and dysfunction.

## Introduction

Mitochondria are essential eukaryotic organelles that generate energy for the cell through oxidative phosphorylation within the respiratory chain. Mitochondrial dysfunction has been implicated in numerous diseases such as cancer, aging, inflammation and neurodegenerative diseases, as well as the process of ageing, making the elucidation of their physiological function and dysfunction a subject of significant interest^1–3^. *In situ* characterization of the physical and chemical properties of proteins in a native physiologically relevant environment is crucial for understanding fundamental biological processes and the pathogenesis of these diseases. However, obtaining atomic-level insights into protein structure and interactions within intact mitochondria remains a significant challenge due to methodological limitations.

In-cell NMR spectroscopy is a powerful tool for studying protein behavior in living biological systems, offering direct observations of proteins in their native cellular environment and bridging the gap between *in vitro* structural biology and in vivo cellular biology^4–6^. Over the past decade, significant advancements have been made in protein labeling, delivery, and expression, as well as in NMR sampling strategies^7–11^. These developments have expanded the applicability of in-cell NMR from prokaryotic cell to eukaryotic and even human cells, providing a critical foundation for investigating disease mechanisms and advancing drug development.

Although powerful in principle, in-cell NMR still faces challenges, one major limitation is that overexpressed or delivered proteins are often distributed uniformly throughout the cytoplasm, as observed in A2780 and RCSN-3 cells during in-cell studies^12^. This uniform cytoplasmic localization overlooks subcellular effects that can influence protein interactions. Most eukaryotic proteins perform their physiological functions within specific cellular compartments, and some even exhibit distinct functions depending on their subcellular localization^13^. Consequently, whole-cell level studies may lack physiological relevance and, in some cases, lead to misinterpretations. To address this issue, recent studies have employed signal peptides to direct expressed or delivered proteins to specific organelles for NMR detection, or to study the endogenous protein in isolated organelles, providing insights into protein folding, conformational changes and interactions within these organelles ^14–17^. However, due to the limitations in protein expression levels and localization efficiency, as well as undesired background signals in mammalian expression systems, the atomic insights of proteins in organelles is still very limited.

Here, we present an approach that enables NMR observation of proteins within intact mitochondria by delivering proteins into animal tissue–derived mitochondria via electroporation. The method was applied to a disordered protein, α-synuclein (α-syn), as well as three globular proteins. Using this in-mitochondria NMR technique, we characterized the interactions of α-syn with mitochondria membranes. Additionally, we successfully studied the stability of *streptococcus* protein G B1 domain (GB1) and the dimerization of its variant within intact mitochondria, achieving quantitative determination of GB1 dimerization using a chemically labeled, high-sensitivity tag. Our work provides an effective means for *in situ* studies of proteins in intact mitochondria at atomic level.

## Results

### Delivering proteins into mitochondria via electroporation for *in situ* NMR studies

Mitochondria are abundant in energy-demanding mammalian tissues such as the heart, liver, skeletal muscle and brain. To establish a robust method for *in situ* NMR studies in intact mitochondria, we isolated large quantities of mitochondria from mitochondria-rich tissues, including liver and brain, according to previously reported protocols^18,19^.

Isotope-labelled proteins were produced and purified from *E coli*. and subsequently delivered into mitochondria via electroporation. The protein-loaded mitochondria were then utilized for in-mitochondria NMR detection (**Fig. 1a**). The purity of the isolated mitochondria was confirmed by Western blot analysis, using marker protein antibodies to ensure minimal contamination from other cellular components (**Fig. 1b**). Mitochondrial viability and membrane integrity were assessed using the fluorescent dye Mito-Tracker and the citrate synthase assay (**Fig. 1c,d**). The high red fluorescence signal indicated a strong membrane potential, while the low citrate synthase activity confirmed the integrity of the mitochondrial inner membrane, demonstrating that the mitochondria remained functional and healthy. These results confirm that the protocol produces viable mitochondria and that the electroporation processes have minimal impact on their viability and integrity. The supernatant was collected after the in-mitochondria NMR experiment to assess whether the signals were from the leaked protein during the experiment.

**Fig.1.**
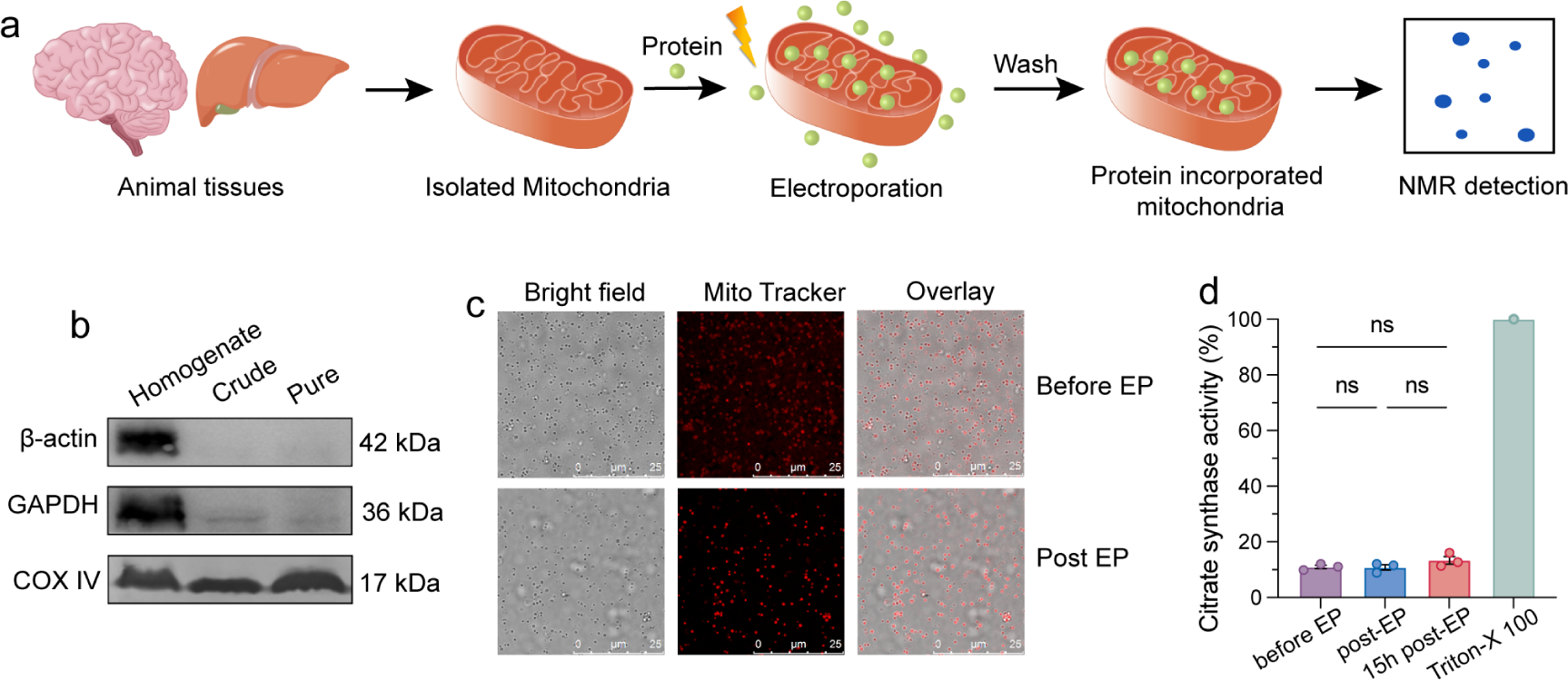
In-mitochondria NMR sample preparation. a) Schematic diagram of in-mitochondria NMR sample preparation. b) Western blot assay of isolated mitochondria using β-actin, GAPDH and Cox IV antibody. c) Fluorescence images of isolated mitochondria stained with Mito Tracker Red CMXRos before and after electroporation. Scale bar, 25 μm. d) Citrate synthase activity assay of isolated mitochondria before and after electroporation. One-way ANOVA was used for comparison of more than two groups (ns: no statistical significance).

### Labeling strategy for *in situ* study on proteins using in-mitochondria NMR

Proteins are produced in *E coli.* with isotope-labeling and then delivered into the mitochondria via electroporation. We first take an intrinsically disordered protein α-synuclein (α-syn) as an example, which has been reported to play a role in the pathology of Parkinson’s Disease (PD)^20,21^. Uniform (U)-^15^N-enriched α-syn was delivered into mitochondria and its two-dimensional (2D) ^1^H-^15^N SOFAST HMQC spectrum was then acquired. Given the high consistency of α-syn spectra in mitochondrial derived from liver and brain (**Extended Data Fig. 1a,b,c,d,e**), liver mitochondria were selected for subsequent experiments due to the ease of extraction and the ability to obtain large sample quantities. The spectrum of α-syn in mitochondria displayed a narrow dispersion in the ^1^H dimension (**Fig. 2a**), consistent with that in buffer, indicating that the mitochondrial environment did not induce a major conformational rearrangement of α-syn. We observed line broadening and overlapping of resonances in the in-mitochondria NMR spectra, mainly due to the viscosity and interactions of α-syn with endogenous cellular components. To overcome the signal overlap and to better resolve the in-mitochondria spectra, specific labeling was then employed to reduce the number of observable peaks. We prepared either Lysine/Tyrosine-selective or Methionine-selective ^15^N-enriched α-syn and collected its in-mitochondrial ^1^H-^15^N correlation spectrum to complement the resonances that were failed to be identified in the spectrum of the U-^15^N-enriched protein due to line-broadening and signal overlapping (**Fig. 2b,c, Extended Data Fig. 1f,g**).

**Fig. 2.**
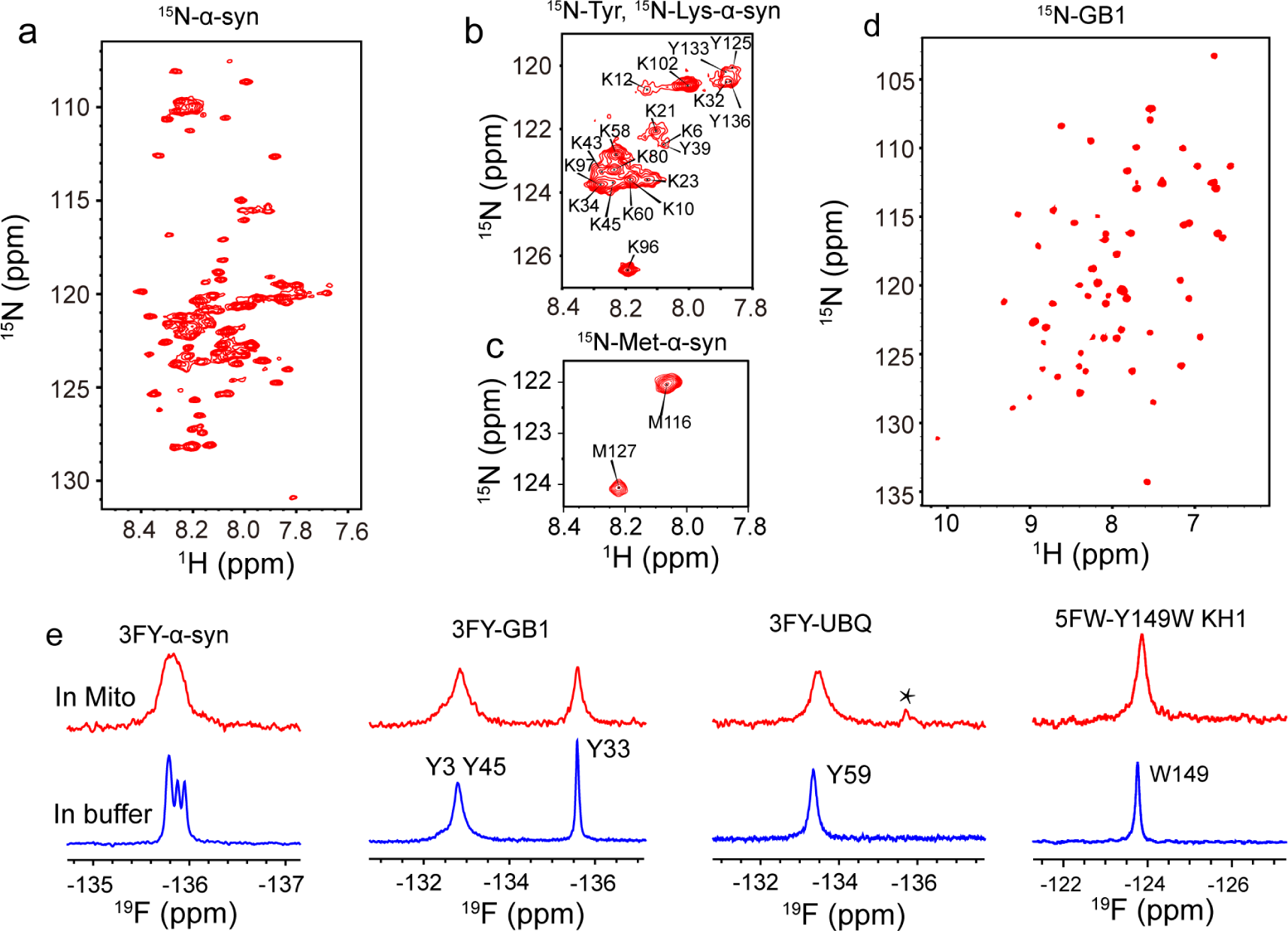
In-mitochondria NMR spectra of proteins with ^15^N enrichment and ^19^F labeling. a-c) ^1^H-^15^N spectra of ^15^N-enriched α-syn (a), Lysine/Tyrosine-selective (b), Methionine-selective ^15^N-enriched α-syn (c) and GB1 (d) in mitochondria. e) ^19^F spectra of 3FY labeled α-syn, GB1, UBQ and 5FW labeled KH1-Y149W in mitochondria.

In contrast to disordered proteins, it is challenging to obtain atomic resolution information for many globular proteins in the cellular environment due to the fact that the interaction of globular proteins with large cellular components often broaden the resonances in conventional heteronuclear NMR spectra severely or even beyond detection, as well as the relatively low delivery efficiency of electroporation for structural proteins compared to disordered proteins. Here, three U-^15^N-enriched globular proteins, GB1, human ubiquitin (UBQ) and a domain of human K-homology splicing regulator protein (KH1) were delivered into mitochondria via electroporation, and corresponding 2D ^1^H-^15^N correlation spectra were collected. As shown in **Fig. 2d**, GB1 yielded a well-resolved in-mitochondria NMR spectrum similar to that of the protein in buffer, showing a pattern typical of a stably folded protein, consistent with previous results in *E coli*^22^. However, the in-mitochondria ^1^H-^15^N correlation spectrum of UBQ showed several overlapping signals that were poorly distributed in the ^1^H dimension (**Extended Data Fig. 2a**). Comparison with the *in vitro* reference spectrum showed that the positions of most of the cross-peaks were not well preserved in the in-mitochondria spectrum, suggesting that the imported UBQ may be involved in interactions with cellular components leading to signal broadening beyond detection, while a fraction of the protein is degraded in mitochondria, yielding resonances with small chemical shift dispersion and narrow half-peak width. Unfortunately, KH1 did not yield a high quality 2D ^1^H-^15^N correlation spectrum in the mitochondria (**Extended Data Fig. 2b**). These results suggest that for globular proteins, ^15^N labeling is not always applicable for protein studies in the cellular environment.

In recent years, ^19^F labeling has been found to be an ideal nucleus for in-cell NMR studies because it is 100% natural abundant, highly sensitive to its local chemical environment and absent from virtually all natural macromolecules^23–25^. We therefore used ^19^F NMR to perform *in situ* mitochondrial studies on α-syn and three globular proteins, GB1, UBQ and KH1. 3-fluorotyrosine (3FY) was incorporated into the proteins during recombinant expression in *E. coli*. The proteins were delivered into mitochondria and one-dimensional (1D) in-mitochondria ^19^F NMR spectra were acquired. As shown in **Fig. 2e**, line broadening of the ^19^F signals was observed in mitochondria compared to that in buffer. α-syn contains four tyrosine residues, one (Y39) in the N-terminus and the other three (Y125, Y133, Y136) in the C-terminus. The ^19^F in-mitochondria spectrum of 3FY-labelled α-syn showed a single broad resonance where the resonances for four tyrosine residues were broadened into one peak, suggesting that both the N- and C-terminus of α-syn are involved in interactions with large cellular components, which is consistent with the ^15^N-enriched NMR results. GB1 contains three tyrosines, the resonances of Y3 and Y45 overlapped in mitochondria, consistent with the spectrum in buffer. Both UBQ and KH1 contain only one tyrosine, and a single ^19^F resonance was observed in the ^19^F 1D spectrum of 3FY labeled UBQ and 5FW labeled KH1-Y149W, respectively. The successful acquisition of in-mitochondria NMR spectra provides a means to study protein structure and functional mechanism in intact mitochondria.

### α-syn is involved in membrane interactions in mitochondria via its N-terminus

There is growing evidence that the binding of α-syn to mitochondrial membranes is pathologically important^26,27^, but it is still remains a challenge to observe the interactions *in situ* at the atomic level. The in-mitochondria 2D ^1^H-^15^N HMQC spectrum of α-syn exhibited varying degrees of NMR signal line broadening for the N-and C-terminal residues (**Fig. 3a**), suggesting that the interactions involve these regions in mitochondria. The residue-resolved signal attenuation profile of α-syn N-terminus in mitochondria exhibits the *in-vitro* membrane interaction signature ^28^, where the large size of membranes broaden the resonances of the corresponding amide group upon binding of the residues to membranes, we therefore speculated that the delivered α-syn is likely to be involved in membrane interactions in mitochondria. Further *in-vitro* characterization of α-syn in the presence of the membrane fraction isolated from mitochondria was performed. It is found that the signal from the N-terminus of α-syn is severely attenuated in the presence of mitochondrial membrane (**Fig. 3b**), which is highly consistent with the profile of α-syn in the presence of mitochondrial membrane-mimicking liposomes containing the mitochondria-specific phospholipid, cardiolipin (CL) (**Extended Data Fig. 3a**). To further confirm the interactions of α-syn with mitochondrial membranes, the PD-related familial mutant A30P, which has been reported to have a reduced binding affinity to synaptic membranes ^29^ was delivered into mitochondria, and a decrease in the attenuation of the NMR resonances for the N-terminal residues was observed compared to that of WT α-syn (**Fig. 3c**), indicating a reduced interaction of the A30P mutant in mitochondria, which was highly consistent with the results of the in vitro titration experiment of A30P with mitochondrial membrane-mimicking liposomes (**Extended Data Fig. 3b**). Taken together, these findings suggest that the N-terminus of α-syn is mainly involved in the interaction with the membranes in mitochondria. We also observed signal attenuation of the C-terminal residues in mitochondria, which indicate that the C-terminus is involved in interactions in mitochondria. Further studies on exploring the binding partners of the C-terminus may provide valuable insights into the biological and pathological role of α-syn in mitochondria.

**Fig. 3.**
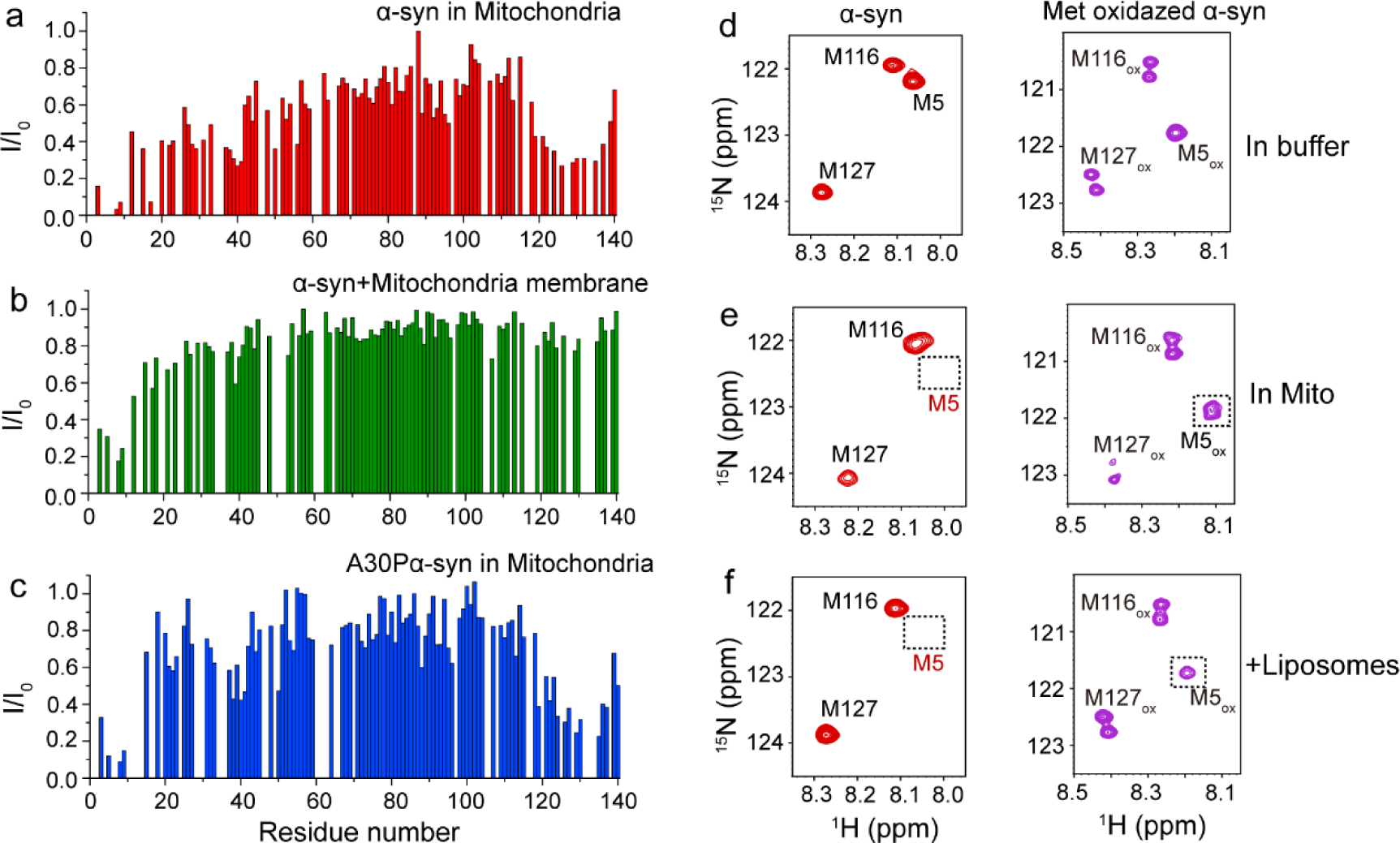
α-syn involves in the membrane interactions in mitochondria. (a-c) Residue-resolved signal attenuation of ^15^N-enriched α-syn in mitochondria (a), in the presence of mitochondrial membrane (b) or its A30P mutant in mitochondria (c) compared to that in buffer. (d-f) ^15^N-enriched reduced (left) or oxidized (right) α-syn in buffer (d), mitochondria (e) or in the presence of liposomes (f).

Cellular oxidative stress is one of a cascade of factors contributing to the degeneration of dopaminergic neurons in PD, with an imbalance of reactive oxygen species (ROS) leading to oxidative modification of α-syn on Methione (Met)^30,31^. Here, we employed In-mitochondria NMR to explore whether oxidation affects the interactions of α-syn involved in mitochondria. α-syn was Met-selectively ^15^N-labeled to simplify the in-mitochondria spectrum and facilitate the identification of each Met with the reduced and oxidized states, respectively. Met oxidation was achieved by treating the protein with 4% H2O2, and complete oxidation of Met5, Met116 and Met127 was verified by recording ^1^H-^15^N HMQC spectra (**Fig. 3d, Extended Data Fig. 4a**). Oxidized and reduced Met-selective ^15^N-enriched α-syn were respectively delivered into mitochondria and the corresponding ^1^H-^15^N HMQC spectra were collected. As shown in **Fig. 3e**, the 2D NMR spectra of reduced Met-^15^N α-syn in mitochondria revealed only three well-resolved amide resonance cross peaks for Met116 and Met127, in which the resonance of Met5 was broadened beyond detection, probably due to the strong interaction of the N-terminus in mitochondria. Notably, the cross-peak of Met5 from oxidized α-syn remained intact in mitochondria, likely because the interaction of α-syn on Met5 in mitochondria is disrupted by oxidation. The role of oxidation in modulating α-syn’s membrane interaction was further supported by *in vitro* NMR titration experiments, which show that the interaction of α-syn with liposomes is abolished by oxidation (**Fig. 3f**), consistent with the results of a previous study that oxidation of α-syn methionine reduces its affinity for biological membranes^32^. Additionally, NMR titration results revealed that α-syn has minimal interaction with the mitochondrial chaperone tumor necrosis factor receptor-associated protein 1 (TRAP1), but interacts with protein disulfide-isomerase (PDI), with this interaction being regulated by Met oxidation (**Extended Data Fig. 4b-e**). Together, these findings suggest that α-syn is involved in membrane interactions in mitochondria via its N-terminus and that these interactions are regulated by Met oxidation.

### Characterizing protein folding and stability in mitochondria

Protein stability is determined by the delicate balance of numerous attractive and repulsive interactions that govern whether proteins fold or unfold, as well as whether they associate or dissociate with their binding partners. NMR is a powerful technique for quantifying the kinetics and equilibrium thermodynamic stability of structured proteins and their complexes^33,34^. While cytoplasm is known to influence protein stability in cells, the measurement of the subcellular microenvironment on protein stability is still lacking. Here, the effect of the mitochondrial environment on folding stability of GB1 was monitored by time-resolved ^1^H-^15^N HSQC spectra, as described in literature^35^. Simply put, the signal recover rate of deuterated protein indicates relative stability, with more stabilized protein exhibiting slower recover rate.

The ^1^H-^15^N HSQC spectra of ^15^N-enriched, deuterated GB1 were recorded at identical time points in both buffer and mitochondria, respectively. GB1 produced well-resolved 2D spectra in mitochondria, but some cross-peaks were missing compared to the *in vitro* reference spectrum (**Fig. 4a,b, Extended Data Fig. 5**), suggesting that certain backbone amide deuterium was not effectively exchanged for some residues in mitochondria. A detailed analysis of time-resolved signal recovery of the deuterated protein in buffer and mitochondria is shown in **Fig. 4c,d**. In solution, residues in the loop region showed rapid signal recovery, while those at the protein core recovered more gradually. In mitochondria, however, residues in both the loop and the structural regions exhibited slower signal recovery. With many residues not fully exchanged throughout the 17-hour sampling period. This reduced exchange rate in mitochondria is likely due to non-specific interactions and crowding effects within the mitochondria environment, which contribute to enhanced stability of GB1 in mitochondria compared to that in buffer.

**Fig. 4.**
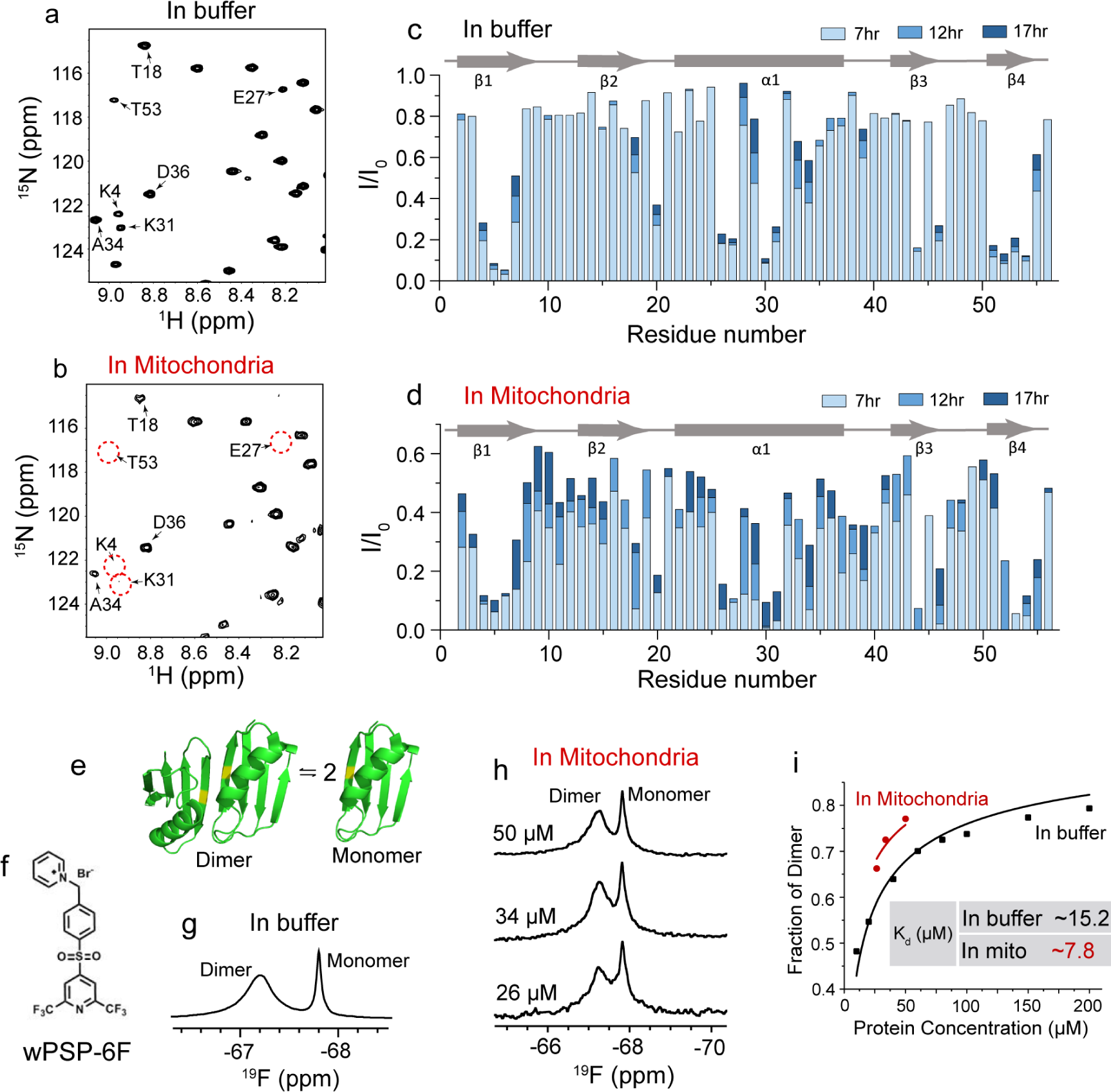
Protein stability measurement in mitochondria. (a-b) ^1^H-^15^N HSQC spectra of ^15^N-enriched deuterated GB1 in PBS buffer (pH 6.9, 288K) (a) and mitochondria (b). (c-d) Backbone amide hydrogen-deuterium exchange of GB1 in buffer and in mitochondria. Residue-resolved signal attenuation of ^15^N-enriched deuterated GB1 in buffer or mitochondria at different time compared to ^15^N-enriched GB1 (I/I0) were shown. (e) Dissociation of the A34F GB1 side-by-side homodimer (PDB ID code 2RMM) showing the mutation site of T11C where the wPSP-6F tag (f) modified. (g-i) ^19^F NMR spectrum of wPSP-6F tagged A34F/K10N/T16C GB1 variant in buffer (g) and in mitochondria (h), and corresponding fitting curves of dissociation constant (i).

Protein-protein interactions are fundamental to signaling transduction and various cellular processes. Acquiring quantitative equilibrium data on protein complexes within cells is essential for understanding their functions. While recent advances have enabled equilibrium thermodynamic and kinetic studies of protein-protein interactions in living cells^9,34^, quantification of these interactions within specific subcellular microenvironment remains limited.

To investigate protein-protein interactions in mitochondria and examine the effects of the mitochondrial microenvironment, we used A34F GB1, a protein that exists in a monomer-dimer equilibrium in buffer (**Fig. 4e**), as a model for binding affinity quantification. We characterized A34F GB1 dimerization using ^19^F NMR as described^34^. However, the ^19^F resonance of 3FY-labelled GB1 A34F exhibited temperature-dependent broadening, with significant overlap between the monomer and dimer resonance of Y3. This poor resolution, coupled with the relatively low sensitivity of 3FY labeling for quantifying binding affinity at low protein concentrations, making us choose the highly sensitive probe, wPSP6F^9^, containing two trifluoromethyl groups to measure the dimerization of A34F GB1 in mitochondria (**Fig. 4f, Extended Data Fig. 6)**. A34F GB1 was chemically labelled at Cys16 with wPSP6F, enabling clear resolution of monomer and dimer populations in mitochondria, and the population of GB1 dimer was determined by integrating the ^19^F resonances as a function of protein concentration (**Fig. 4g-i**). The binding isotherms were then fitted to calculate the equilibrium dissociation constant, *Kd*. Our results revealed that A34F GB1 dimerization exhibited a twofold increase in binding affinity within mitochondria compared to buffer. Together with previous studies in different cellular environments^9,34^, these findings suggest that the A34F GB1 homodimer is more stable within the cellular environment than in buffer, with stability varying across different microenvironments.

## Discussion

In eukaryotic cells, proteins perform their functions within specific subcellular compartments, and some proteins translocate to different subcellular compartments in response to cell signaling^13^. Thus, studying proteins in their native microenvironment with subcellular resolution is highly desirable. In this study, we demonstrate that proteins can be efficiently delivered into isolated mitochondria derived from animal tissues via electroporation, thus providing sufficient samples for NMR experiments and enabling atomic-resolution insights into proteins in intact mitochondria. To our knowledge, this is the first *in-situ* NMR application where proteins are directly delivered into organelles via electroporation.

A key prerequisite for biomolecular NMR applications is that the molecules contain suitable NMR-active isotopes. In-cell NMR studies of macromolecules typically rely on isotopic labeling with ^15^N, ^13^C and ^19^F nuclei. Direct delivery of purified proteins into cells via electroporation allows proteins to be pre-labelled by a variety of labeling strategies, including biosynthetic incorporation, chemical tagging, and covalent modification, which enable the study of post-translational modifications. This approach potentially allowing more proteins to be visualized and more molecular details to be obtained within the *in situ* environment. Moreover, It ensures that only the target protein is labelled, avoiding background signals from cellular components—a common issue in *in situ* expression systems that require relatively high protein expression levels. In this work, we deliver oxidized ^15^N-Met selectively labelled α-syn into mitochondria to examine, at the atomic level, the effect of Met oxidation on α-syn’s interaction with mitochondrial membranes. To explore protein complex stability and quantify binding affinity within the cellular environment, high resolution, high signal-to-noise NMR spectra are essential. However, obtaining satisfactory NMR spectra at low protein concentrations within the cellular milieu is extremely challenging for traditional NMR labeling strategies. By using electroporation to deliver GB1 A34F protein, labelled with the high-sensitivity chemical tag, into mitochondria, we are able to record NMR spectra at low protein concentrations and quantitatively evaluate binding affinities in intact mitochondria.

Unlike prokaryotic cells, eukaryotic cells are compartmentalized by membranes that segregate biochemical reactions into distinct organelles. The cellular microenvironment, composed of dynamically interacting entities, varies significantly across different organelles. Proteins interact with their surroundings to carry out biological functions within these specific cellular environments. Intracellular seeding events of α-syn have been shown to preferentially occur on mitochondria membrane surface^27^. Our study demonstrates that α-syn primarily interacts with the membranes in mitochondria, in contrast to its interactions in the cytoplasm, where it has been reported to interact with chaperones^36^. A defining feature of the mitochondrial membrane is the presence of CL, a unique phospholipid not found in the cytoplasm^37^. Previous studies have shown that α-syn readily interacts with CL membranes^26^, which likely explain its distinct interaction patterns in different environments. These findings highlight the importance of studying proteins in specific subcellular environments to better understand their physiological behavior and interactions. Our in-mitochondria NMR approach offers a promising tool for obtaining atomic-level insights into the structure and function of proteins involved in mitochondrial-specific biological processes, such as mitochondrial protein import, oxidative phosphorylation and the expression of mitochondrial genes.

In summary, we present a powerful approach for in-organelle NMR studies by delivering proteins directly into animal tissue-derived mitochondria via electroporation, providing valuable insights into protein interactions and stability in intact mitochondria. The methodology is readily transferable to other organelles or those isolated from animal disease models, offering a powerful tool for investigating the pathogenesis of organelle-related diseases and exploring potential therapeutic strategies.

## Methods

### Protein preparation

The plasmids containing the coding sequence for α-syn, α-syn-A30P, GB1, GB1-A34F-K10N-T16C, UBQ and KH1-Y149W were transformed into *Escherichia coli* BL21 (DE3) cells. Single-point mutants were generated by site-directed mutagenesis. ^15^N enriched proteins were produced in M9 minimal media supplemented with ^15^NH4Cl as sole nitrogen resource, and amino acid selective labeling were achieved in M9 minimal media supplemented with ^15^N-methionine/lysine/tyrosine respectively. 3FY and 5FW labeling was achieved by using the fluoro-labeled amino acids and glyphosate. Plasmid-containing cells were grown at 37°C to an optical density at 600 nm (OD600) of 0.6-0.8 and induced with 1 mM IPTG for 6 h before harvest.

The detailed purification procedures have been described previously^22,28^. For α-syn and its variant, the cell lysates were boiling for 20 min, streptomycin sulfate (10 mg/mL) was added to α-syn-enriched soluble fraction to precipitate nuclear acids prior to precipitating α-syn with ammonium sulfate (360 mg/mL). The pellet was resuspended and loaded onto an anion exchange (DEAE Fast Flow column, GE Healthcare) using 20 mM Tris, pH 7.6 with a 0-1 M NaCl gradient. Methionine oxidation was achieved by incubating α-syn monomer with 4% H2O2 for 2 h at room temperature, while excessive H2O2 was removed by multistep ultrafiltration through 10-kDa cutoff membrane (Amicon Ultra). GB1 and its variant were purified using an anion exchange (DEAE Fast Flow column, GE Healthcare) equilibrated with 50 mM Tris, pH 8.0 and eluted by a linear gradient of 0-1 M NaCl. For UBQ, cells were homogenized with 1:1 chloroform/lysis buffer (50 mM sodium acetate, pH 5.0), centrifuged at 10,000 rpm for 30 min to recover the aqueous layer, finally purified by a cation exchange (SP Fast Flow column, GE Healthcare) using 50 mM sodium acetate, pH 5.0 with a 0-1 M NaCl gradient, and a size exclusion chromatography (HiLoad superdex75 16/600 column, GE Healthcare) equilibrated with 50 mM sodium acetate, 250 mM NaCl, pH 5.0. PDI was firstly eluted with a linear gradient of 0-500 mM imidazole on a HisTrap Ni-NTA affinity column (GE Healthcare), followed by an anion exchange (Resource Q column, GE Healthcare) using 20 mM Tris, pH 8.0 with a 0-1 M NaCl gradient, and further purified with a size exclusion chromatography (HiLoad superdex200 16/600 column, GE Healthcare) equilibrated with 20 mM Tris, 300 mM NaCl, pH 8.0. TRAP1 was purified by a HisTrap Ni-NTA affinity column (GE Healthcare) using 20 mM PB, 500 mM NaCl, 10 mM β-mercaptoethanol (β-ME), 20 mM imidazole, pH 7.5 with a 20-500 mM imidazole gradient, the proteins were then concentrated and chromatographed on a Superdex200 16/600 column (GE Healthcare) equilibrated with 20 mM PB, 250 mM NaCl, pH 7.5. Protein concentration was determined by absorbance of aromatic amino acids at 280 nm with a NanoDrop spectrophotometer, lyophilized upon aliquots and stored at −80 °C.

### Chemical tagging of wPSP-6F

The reaction was performed as described^9^. Briefly, 1 equiv of the purified protein was mixed with 3 equiv of wPSP-6F tag and 2-3 equiv of tris (2-carboxyethyl) phosphine (TCEP) in 50 mM Tris (pH 8.0), and shaken at room temperature or 37 °C for 12 h to ensure the high ligation efficiency of over 95%. The tagged protein then was purified by sequential dialysis. The purity and concentration were further determined by ^19^F NMR (internal standard: 4-F phenylalanine).

### Mitochondria isolation

All the procedures for mitochondria isolation were performed on ice. Mitochondria from mice liver were isolated following a method published previously^18^. Briefly, 8 g of liver tissues freshly extracted from 4-week rat were homogenized in cold isolation buffer (10 mM Tris, 1mM EDTA·2K, 320 mM Sucrose, pH 7.4). The homogenate was centrifuged four times at 1,300 *g* for 15 min to remove tissue debris and heavy membranes, then the resulting supernatant was centrifuged at 7,000 *g* for 15 min to pellet mitochondria, followed by washing twice with cold isolation buffer. Pig brains were obtained from a slaughterhouse. Mitochondria from pig brain were isolated following a method published previously^19^. Briefly, the tissue was homogenized and then centrifuged at 1,300g for 10 min to remove cell debris and nucleus. The supernatant was collected and centrifuged at 21,000g for 10min, and the pellet was collected and resuspended in 15% Percoll solution, then carefully layered on the top of freshly made Percoll gradient (23% and 40% Percoll) and centrifuged at 30,700 g for 10 min. The mitochondria-enriched fraction at the interface between the 23% and 40% Percoll layers was then collected, further diluted with isolation buffer and centrifuged at 16,700g for 10 min. The pellet of mitochondria obtained was washed in isolation buffer followed by centrifugation at 7,000g for 10 min.

### Preparation of in-mitochondria samples

A highly purified proportion mitochondria were used to prepare in-mitochondria NMR samples. Isotope-labeled proteins was dissolved in 1 mL of 330 mM Sucrose to a final concentration of 1-5 mM and mixed with mitochondria pellet. The mixture were then transferred into 0.1 cm gap cuvettes and electroporated by one pulse at 1200 V on an Eppendorf Eporator. Samples post eletroporation were transferred into cold wash buffer (137 mM KCl, 10 mM Tris, 0.5 mM EDTA·2K, 2.5 mM MgCl2, pH 7.4) and washed for three times by centrifugation at 7,000 *g* for 15 min.

### Confocal microscopy

The isolated mitochondria were stained with 250 nM MitoTracker Red CMXRos for 30 min and then washed with isolation buffer for three times to remove any residual dye. The mitochondria were observed under a fluorescence confocal microscope (Leica TCS Sp8, Germany). The excitation and emission wavelength are 579 nm and 599 nm, respectively.

### Western blot

Samples were applied to 15% SDS/PAGE and then transferred to PVDF membrane by a Bio-Rad semi-dry transfer system. The membranes were then blocked by 5% skimmed milk powder in TBST buffer (10 mM Tris, 150 mM NaCl, 0.01% Tween 20, pH 8.0) for 1h and incubated with the primary antibody for 2 h at room temperature. The membrane was washed with TBST buffer, incubated with anti-rabbit IgG peroxidase secondary antibody, washed again with TBST buffer for three times, and visualized with the enhanced HPR-DAB chromogenic kit (Qiagen).

### Separation of mitochondrial membrane and matrix

Mitochondria isolated from rat liver were suspended in ultrapure water, and subjected to sonication on ice for 15 min to generate lysates. Mitochondrial lysates were then centrifuged at 150,000 *g* for 60 min at 4 °C to separate mitochondrial membrane (pellet) and matrix content (supernatant). The pellet was resuspended in PBS (20 mM PB, 150 mM NaCl, pH 7.2) buffer and sonicated for several cycles.

### Preparation of liposomes

Liposomes mimicking mitochondrial inner membrane were prepared using POPC and CL from Avanti Polar Lipids as described^1^. Briefly, lipid powders were mixed at the desired ratio and dissolved in chloroform. Solution of lipid mixtures were dried, and lipids were rehydrated with PBS buffer to a final concentration of 24 mM. The lipids finally were sonicated to obtain liposomes.

### NMR experiments

Experiments were performed on Bruker 700 and 850 MHz spectrometers equipped with a HFCN and triple-resonance HCN cryoprobe, respectively. Two dimensional (2D) ^1^H–^15^N spectra were acquired with spectral widths of 12 (^1^H) and 26 ppm (^15^N), 0.1 s relaxation delay, 2048 complex points along t1 dimension and 200 in t2 dimension. Spectra of α-syn were typically collected at 279K, whereas spectra for GB1, UBQ, and KH1-Y149W were collected at 288K. The data were processed with Topspin and NMRPipe, and analyzed with SPARKY.

### Citrate synthase assay

Mitochondrial inner membrane integrity was assessed by measuring citrate synthase activity in isolated mitochondria before and after membrane disruption by the detergent Triton X-100 (0.2%) using the citrate synthase assay kit, the formation of thionitrobenzoic acid (TNB) was tested by the absorbance at 412nm.

### Intramitochondrial pH measurement

BCECF was loaded into isolated mitochondria as an indicator of pH in matrix by incubating 5 μM BCECF-AM with mitochondria for 1 h at 4 °C, unabsorbed BCECF-AM was removed by washing twice with isolation buffer. For the calibration of pH in mitochondria, the emission at 530 nm was collected in a pH range of 6.6 to 8.8 in the presence of 5 μM CCCP, a protonophore that allows to equilibrate the pH between buffer and the mitochondria matrix, when fluorescence from BCECF was excited at 439 nm and 490 nm respectively, and the fluorescence ratio (F490/F439) could be converted to pH by equation F490/F439 = (A + B × 10(7–pH))/(C + 10(7–pH)) fitted from least-square by which we determined the pH in mitochondria, where A is 1.025, B is 2.372 and C is 0.2235^38^.

## Acknowledgments

This work is supported by grants from the National Natural Science Foundation of China (21925406, 22274161, 21991080), the Ministry of Science and Technology of China (2021YFA1302600) and the Chinese Academy of Sciences (XDB0540000, YSBR068).

## Author contributions

Z.Z., C.Z., G.X., R.D. and J.Y. performed in-mitochondria NMR sample preparation. Z.Z. and C.Z. performed NMR experiments. X.L. provided PDI and TRAP1 protein. Z.C. provided the highly sensitive probe, wPSP6F. Q.W. performed fluorescence imaging. Z.Z., C.Z., L.J., M.L. and C.L. analyzed data. Z.Z. and C.L. wrote the manuscript.

---

**Extended Data Fig.1.**
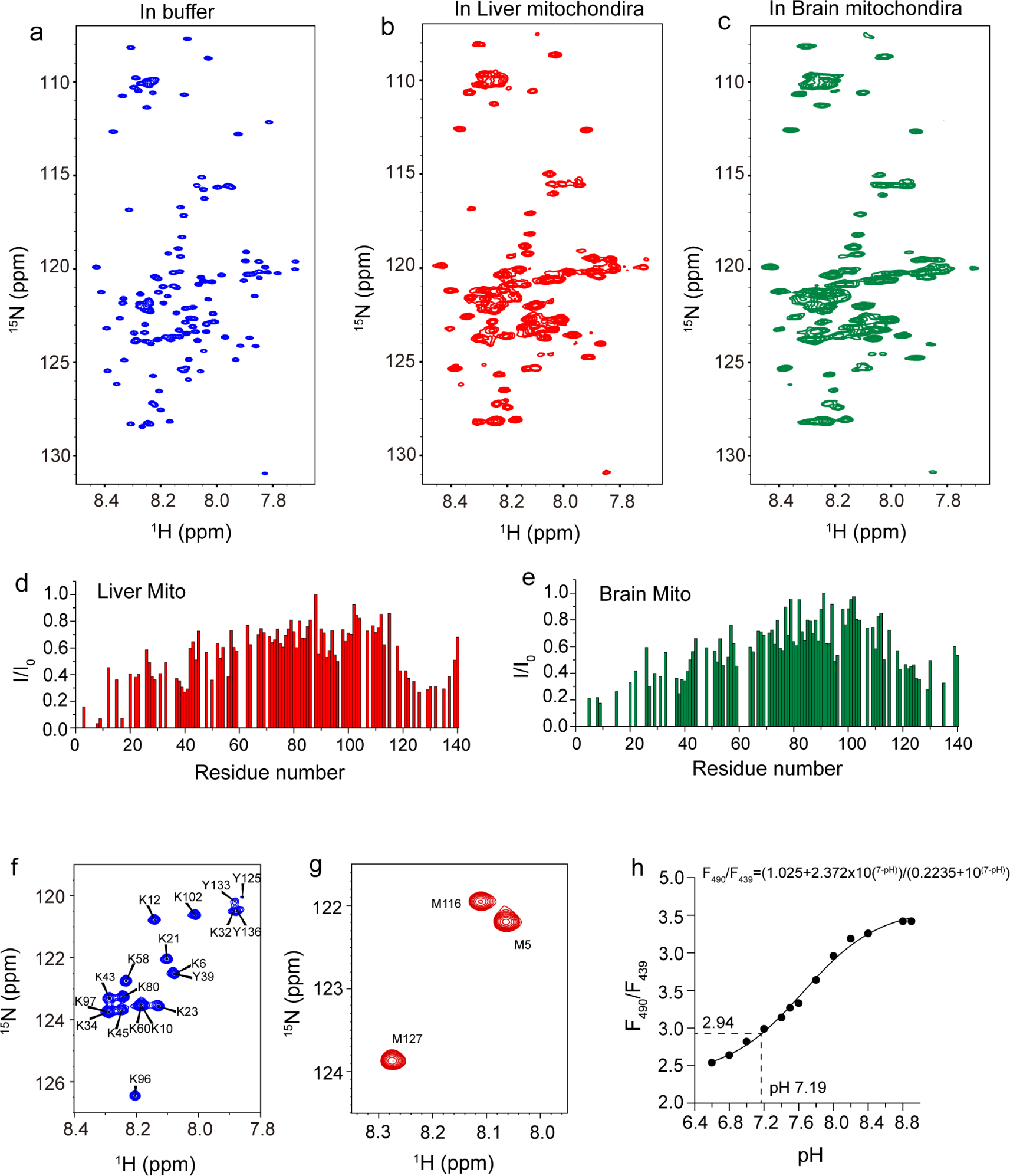
2D ^1^H-^15^N spectra of ^15^N-enriched α-syn in buffer and mitochondria. (a-c) 2D ^1^H-^15^N spectra of uniform ^15^N-enriched α-syn in buffer (a) and mitochondria derived from mice liver (b) or brain (c). (d-e) residue resolved signal attenuation (I/I_0_) of ^15^N-enriched α-syn in mitochondria derived from mice liver (d) or brain (e) compared to that in buffer. (f-g) 2D ^1^H-^15^N spectra Lysine/Tyrosine-selective (f) and Methionine-selective (g) ^15^N-enriched α-syn in buffer. NMR data were collected at 279K. (h) BCECF was loaded into isolated mitochondria to calibrate the pH in mitochondria.

**Extended Data Fig.2.**
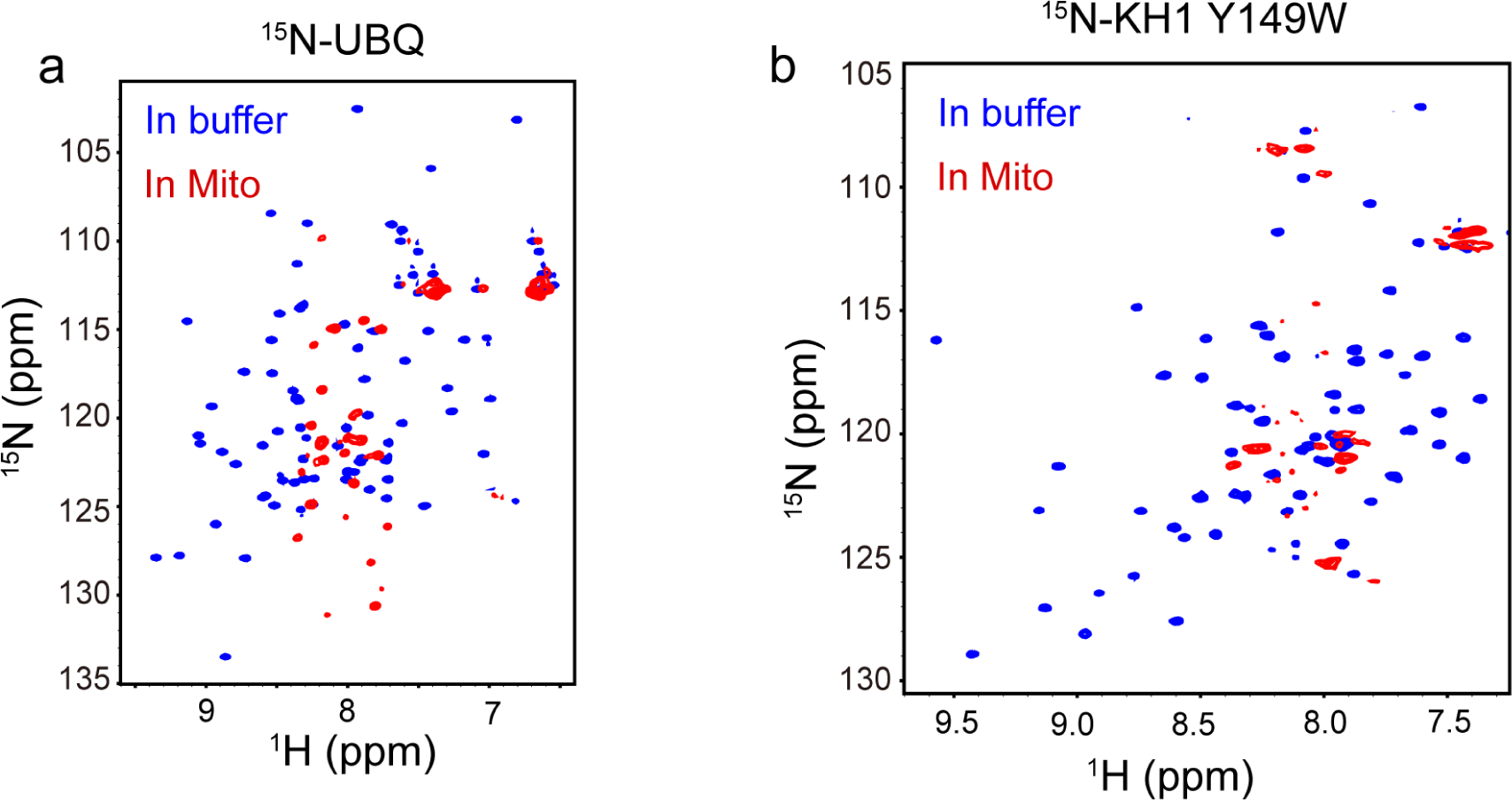
Overlaid 2D ^1^H-^15^N spectra of Uniform ^15^N-enriched UBQ (a) and KH1-Y149W (b) in buffer (blue) and mitochondria (red). NMR data were collected at 288K.

**Extended Data Fig.3.**
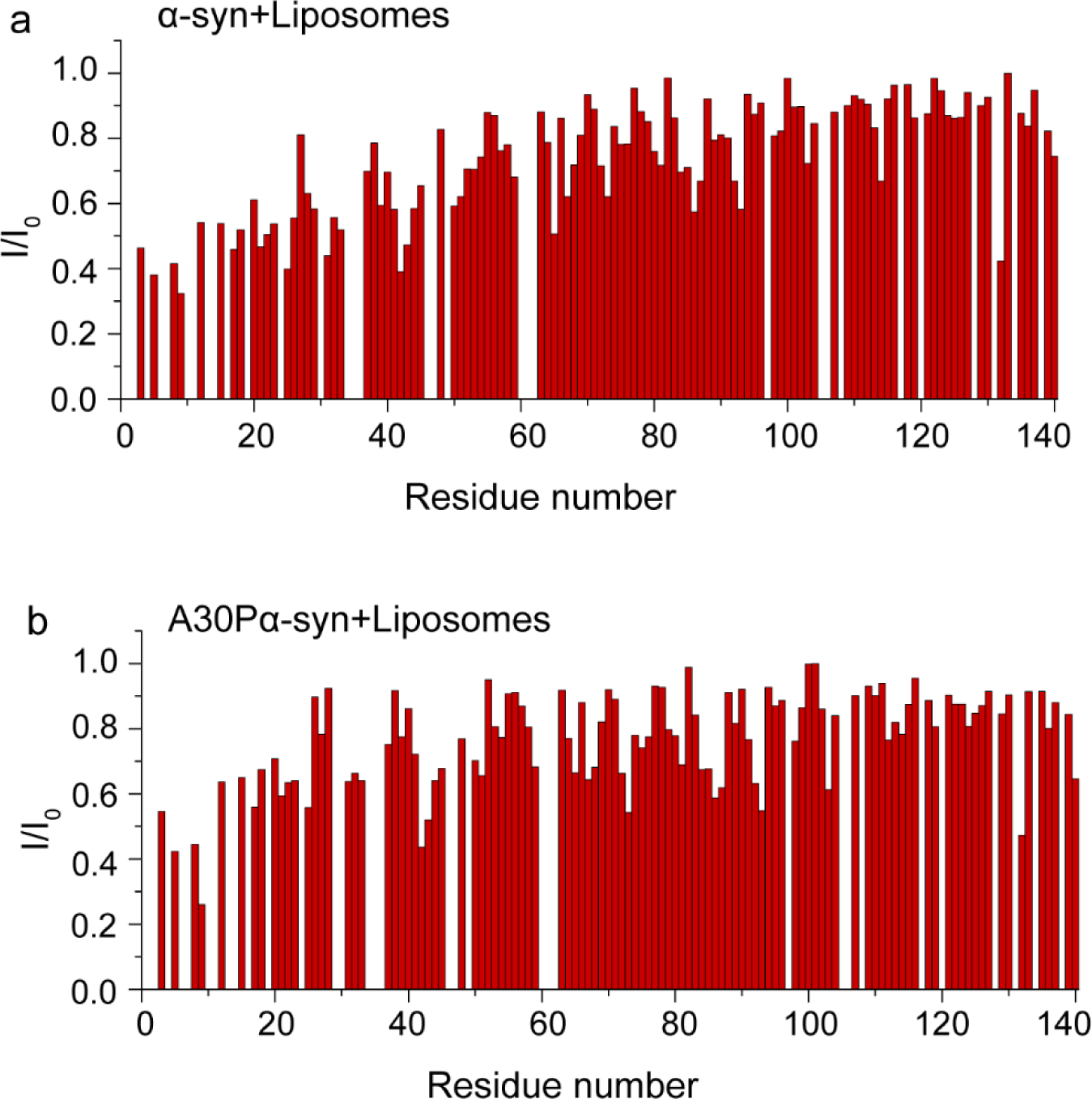
Residue-resolved signal attenuation of ^15^N-enriched α-syn (a) or A30Pα-syn (b) in the presence of mitochondrial membrane-mimicking liposomes compared to that in buffer.

**Extended Data Fig.4.**
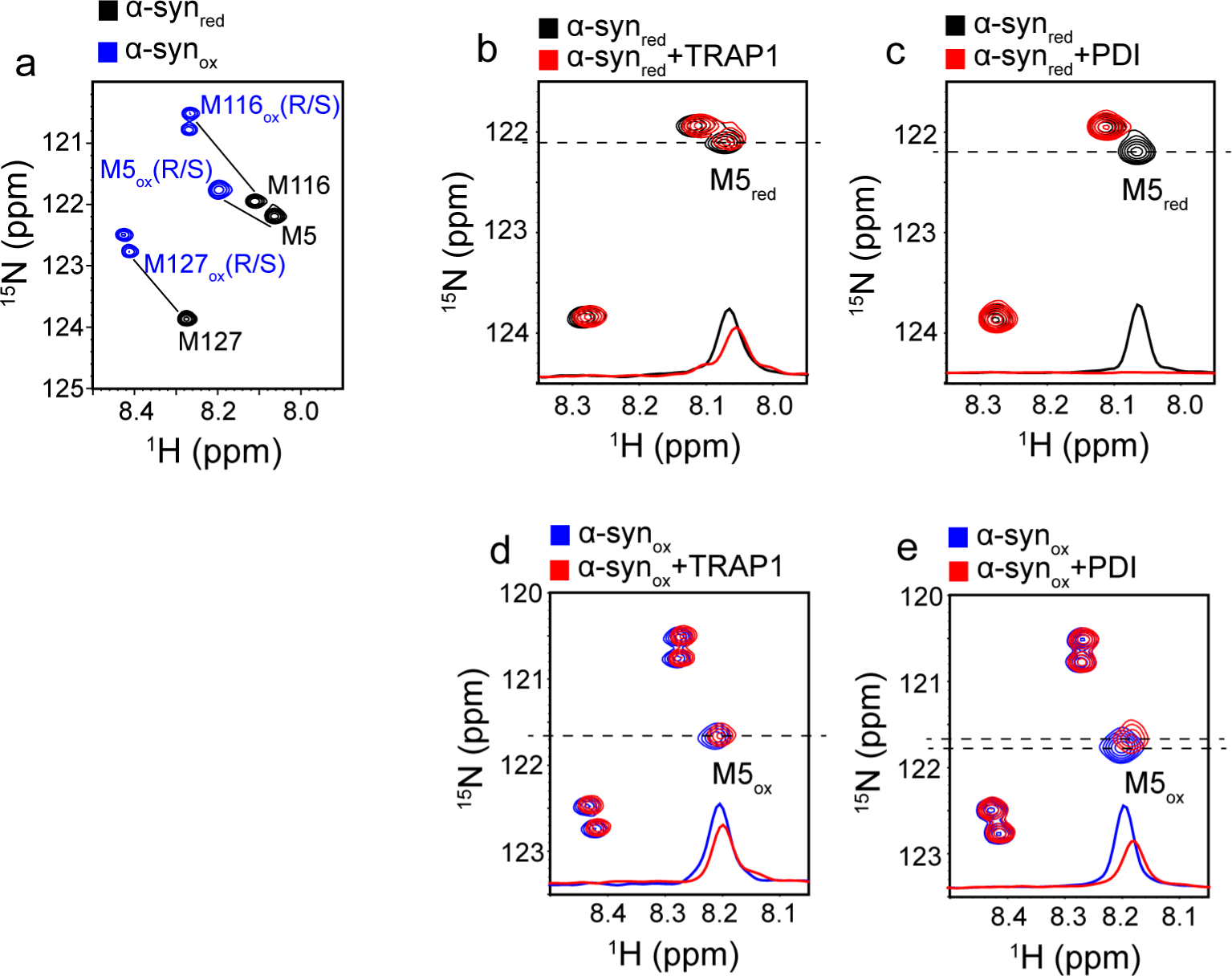
The effect of Met oxidation on the interaction of α-syn with chaperone. 2D ^1^H-^15^N spectra of oxidized (blue) and reduced (black) Met-selective ^15^N-enriched α-syn in buffer (a), in the presence of TRAP1 (b and d) or PDI (c and e).

**Extended Data Fig.5.**
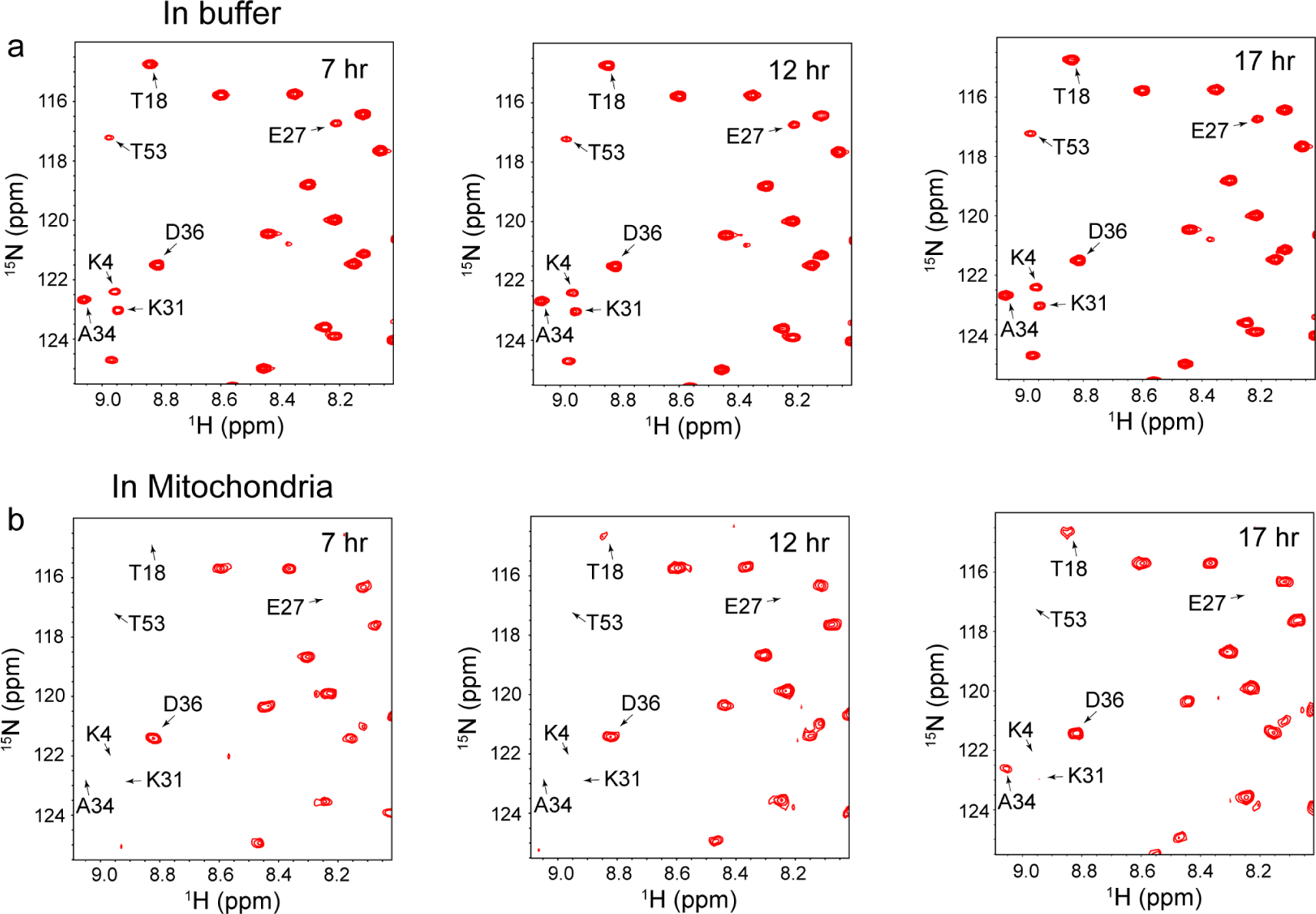
Time-dependent 2D ^1^H-^15^N spectra of ^15^N-enriched deuterated GB1 dissolved in buffer (a) or delivered into mitochondria (b).

**Extended Data Fig.6.**
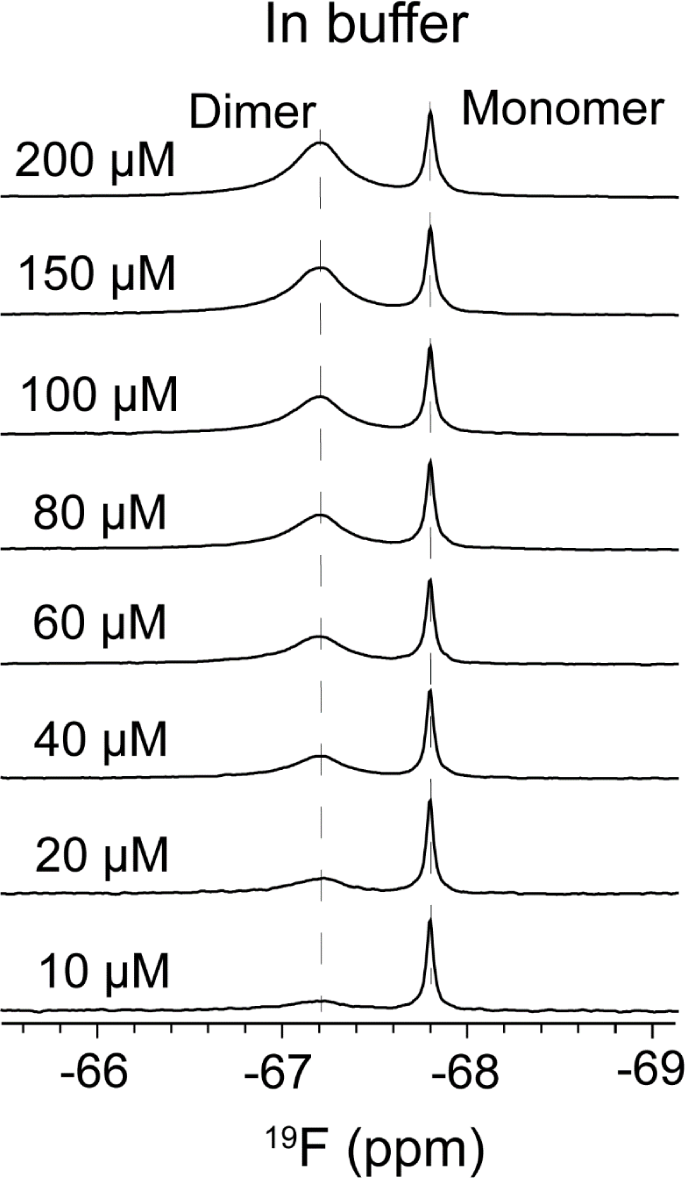
^19^F NMR spectrum of wPSP-6F tagged A34F/K10N/T16C GB1 variant at different concentrations in buffer. NMR data were collected at 288K.

